# Analysis of metabolic pathways in mycobacteria to aid drug-target identification

**DOI:** 10.1101/535856

**Authors:** Bridget P. Bannerman, Sundeep C. Vedithi, Jorge Júlvez, Pedro Torres, Vaishali P. Waman, Asma Munir, Vitor Mendes, Sony Malhotra, Marcin J. Skwark, Stephen G. Oliver, Tom L. Blundell

**Affiliations:** Department of Biochemistry, University of Cambridge, Tennis Court Road, CB21GA, Cambridge, UK; Department of Computer Science and Systems Engineering, University of Zaragoza, María de Luna 3, 50018 Zaragoza, Spain; Department of Biological Sciences, Birkbeck College, University of London, London, UK

## Abstract

Three related mycobacteria are the cause of widespread infections in man and are the focus of intense research and drug-discovery efforts in the face of growing antimicrobial resistance. *Mycobacterium tuberculosis*, the causative agent of tuberculosis, is currently one of the top ten causes of death in the world according to WHO; *M*. *abscessus*, a group of non-tuberculous mycobacteria causes lung infections and other opportunistic infections in humans; and *M*. *leprae*, the causative agent of leprosy, remains endemic in tropical countries. There is an urgent need to design alternatives to conventional treatment strategies, due to the increase in resistance to standard antibacterials. In this study, we present a comparative analysis of chokepoint and essentiality datasets that will provide insight into the development of new treatment regimes. We illustrate the key metabolic pathways shared between these three organisms and identify drug targets with a wide metabolic impact that are common to the three species. We demonstrate that 72% of the chokepoint enzymes are proteins essential to *Mycobacterium tuberculosis*. We show also that 78% of the drug targets, prioritized based on their presence in multiple paths on the metabolic network, are present in pathways shared by *M. tuberculosis, M*. *leprae* and *M*. *abscessus*, including biosynthesis of amino acids, carbohydrates, cell structures, fatty acid and lipid biosynthesis. A further 17% is found in the prioritised pathways shared between *M. tuberculosis* and *M*. *abscessus*. We have performed comparative structure modelling of potential drug targets identified using our analysis in order to assess druggability and demonstrate the importance of chokepoint analysis in terms of drug target identification.

**AUTHOR SUMMARY:** Computer simulation studies to design new drugs against mycobacteria

## Introduction

Pathogenic mycobacteria, such as *M. tuberculosis*, the causative agent of tuberculosis in humans, and *M. leprae,* the causative agent of leprosy, have developed complex metabolic features that enable them to explore their hosts’ environments and compete with the host for nutrients. While *M. leprae* evolved by substantially reducing the size of its genome, losing many metabolic pathways and undergoing further pseudogenization (1,2), *M. tuberculosis* retained metabolic redundancy and an ability to use many different types of sources for nitrogen, carbon and energy, such as sugars, host-derived fatty acids, glucose, tricarboxylic acids, amino acids and cholesterol (3). However, during infection it mostly relies on host-derived fatty acids as the primary energy source (3). *M. abscessus,* a mycobacterium that causes opportunistic infections in cystic fibrosis patients, has developed numerous evolutionary metabolic adaptations (4–6).

Metabolism refers to the chemical reactions organised into pathways and occurring in the cells of all living organisms. These pathways are broadly classified in two main categories: anabolic and catabolic, and are specialised to enable cell survival and replication. Comparative analyses of metabolism in mycobacteria can highlight and identify unique metabolic features, which will be useful in disease management and drug target identification.

Catabolic pathways are energy-producing processes whereby nutrients are broken down into smaller molecules; typical examples are glycolysis, the Krebs cycle, electron transport chain, and fermentation (3,7–10). These are ancient pathways, widely conserved in most organisms, including mycobacteria. Glycolysis and the Krebs cycle, in addition to other energy-producing processes, such as the glyoxylate, pentose phosphate pathway and the electron transport chain, are among the core energy metabolic pathways in *M. tuberculosis, M. abscessus* and *M. leprae* (11,12).

Anabolic pathways are biosynthetic processes comprising the energy-consuming reactions that generate the biomass of mycobacterial cells by synthesizing cellular components such as lipids, carbohydrates, amino acids, and nucleic acids (7,8). These pathways are specialised in mycobacteria and enable them to maintain their structural integrity amidst harsh cellular conditions such as depletion of nutrients and responses of the host immune system during infection. Examples of conserved anabolic pathways across mycobacterial species are the amino acid and nucleic acid biosynthetic pathways. Of particular note is the complex arginine pathway, with a variety of anabolic arginine genes in *M. tuberculosis* (13), the lactic acid bacteria (14) and in *Pseudomonas* (15). L-arginine also occurs as a source of both carbon and nitrogen in bacteria and other microorganisms (15) and the biosynthetic pathway of L-arginine has been confirmed as essential to *M. tuberculosis* (16).

Enzymes of the arginine biosynthetic pathway, including amino-acid acetyltransferase (ArgA) and N-acetyl-gamma-glutamyl-phosphate reductase (ArgC) in *M. tuberculosis,* respectively encoded by the genes, Rv2747 and Rv1652, have previously been identified as high confidence drug targets (17,18). ArgA catalyses the first of eight enzymatic steps in the biosynthesis of arginine from glutamate as well as being involved in the synthesis of ornithine and proline. This involves the activation and reduction of the gamma-carboxyl group in glutamate to produce glutamate-5-semialdehyde. The subsequent steps, catalysed by acetylglutamate kinase (ArgB) and ArgC, involve the phosphorylation of N-acetyl-L-glutamate to N-acetyl-L-glutamate 5-phosphate and its subsequent reduction to N-Acetyl-L-glutamyl 5-phosphate. Acetylornithine aminotransferase (ArgD) catalyses the reaction from N2-acetyl-L-ornithine to N-acetyl-L-glutamate 5-semialdehyde (13).

During the second stage of arginine biosynthesis, ornithine carbamoyltransferase (ArgF) catalyses the formation of citrulline from carbamoyl phosphate, which is in turn catalysed to (N(omega)-L-arginino) succinate by argininosuccinate synthase (ArgG). L-arginine is produced from 2-(N(omega)-L-arginino) succinate in a reaction catalysed by argininosuccinate lyase (ArgH). The intermediate compound, carbamoylphosphate, also provides an alternative to the second stages of L-arginine biosynthesis via the ornithine biosynthetic pathway. Both arginine and ornithine occur as precursors in diverse arginine degradative pathways and in the biosynthesis of polyamines (19–21). Carbamoyl phosphate is also an intermediate in the synthesis of pyrimidines that are the activated precursors of DNA and RNA and a requirement for cell growth (15).

A comprehensive understanding of the enzymes of essential metabolic pathways in mycobacteria, such as the arginine biosynthetic pathway, is necessary for novel drug discovery and elucidating the role of mutations in drug resistance. Currently, crystal structures of some of the enzymes of the arginine pathway of *M. tuberculosis* H37RV exist in the Protein Data Bank (22), including ArgA (5YGE) (23), ArgB (2AP9), ArgC (2nqt, 2i3g, 2i3a) (18) ArgF (2P2G, 2I6U) (18) and ArgJ (3IT4, 3IT6) (24,25) However, experimental structures of ArgD, ArgG and ArgH are lacking for *M. tuberculosis, M. abscessus* and *M. leprae*.

To understand the metabolism of an organism at a system level, it is necessary to identify and reassemble its constitutive parts (metabolites, reactions and uptake processes) which can then be integrated in genomic-scale metabolic networks that taken together can simulate the normal function of a cell (26). These metabolic reconstructions have been used in many different ways from identifying genes essential for growth and thus establishing new drug targets to modelling quantitative drug-dose responses (27,28). A complementary approach to predicting drug targets for mycobacteria is to elucidate the enzymes catalysing reactions that are the sole producers of a specific metabolite or the sole consumers of that metabolite, by querying the annotated genome of an organism, a process referred to as chokepoint analysis (29). Chokepoint enzymes of mycobacterial genomes are critical to bacterial survival and represent potential drug targets from the metabolic networks of the bacteria.

In this study, we show results from comparative pathway and chokepoint analyses of *M. tuberculosis, M. abscessus* and *M. leprae*. We review essential pathways common in the three species and predict potential drug targets for infectious diseases such as leprosy and tuberculosis. We focus on the arginine biosynthetic pathway. For this, we describe comparative modelling procedures to predict the three-dimensional structures of selected enzymes that lack experimental structures, as well as druggability analyses of potential drug targets. We show how the identification and combination of chokepoint reactions within the metabolic networks can indicate essentiality vital for survival of the pathogen. We show how bringing chokepoint analysis of the reactions together with the ligand-binding ability of the enzymes catalysing them is central to the selection of targets for drug discovery.

## Results

### Pathway analyses

For a comparative analysis of pathways shared between *M. tuberculosis* H37Rv, *M. abscessus* and *M. leprae*, we used the newly reconstructed *M. tuberculosis* model iEKVIII (30) and the draft BioCyc pathway databases of *M. abscessus* and *M. leprae*. There were 297 pathways in *M. abscessus*, 212 pathways in *M. tuberculosis* and 135 pathways in *M. leprae* (S1 Table). The larger number of metabolic pathways in *M. abscessus* is consistent with the fact that it is an opportunistic pathogen with a larger genome size of 5MB (31) compared to the relatively smaller genome sizes of the obligate pathogens *M. tuberculosis* (4MB) and *M. leprae* (3.3 MB). There are fewer metabolic pathways in *M. leprae* as it has undergone significant genomic reduction with a small genome size of 3.3 MB and has relatively poor genome annotation (32).

We found 105 pathways in common between the three mycobacteria (*M. tuberculosis, M. abscessus* and *M. leprae*), 152 pathways shared between *M. tuberculosis* and *M. leprae* and 112 pathways shared between *M. abscessus* and *M. leprae*. The differences in the numbers of shared pathways are mainly due to differences in the number of biosynthetic and/or degradative pathways of these organisms. The loss of genes in *M. leprae* due to reductive evolution has resulted in fewer metabolic pathways. The species-specific metabolic pathways identified in *M. tuberculosis* include processes involved in cellulose binding, hydrolase activity, as well as carbohydrate and polysaccharide binding. In the case of *M. abscessus*, species-specific biological processes include an increase in the number of reactions concerned with the biosynthesis of polyamines, thiols and L-citrulline, as well as the number of membrane transport proteins. We analysed these pathways further to identify drug targets common to the three species using the chokepoint analysis algorithm (Fig 1; S1 File).

**Fig 1:**
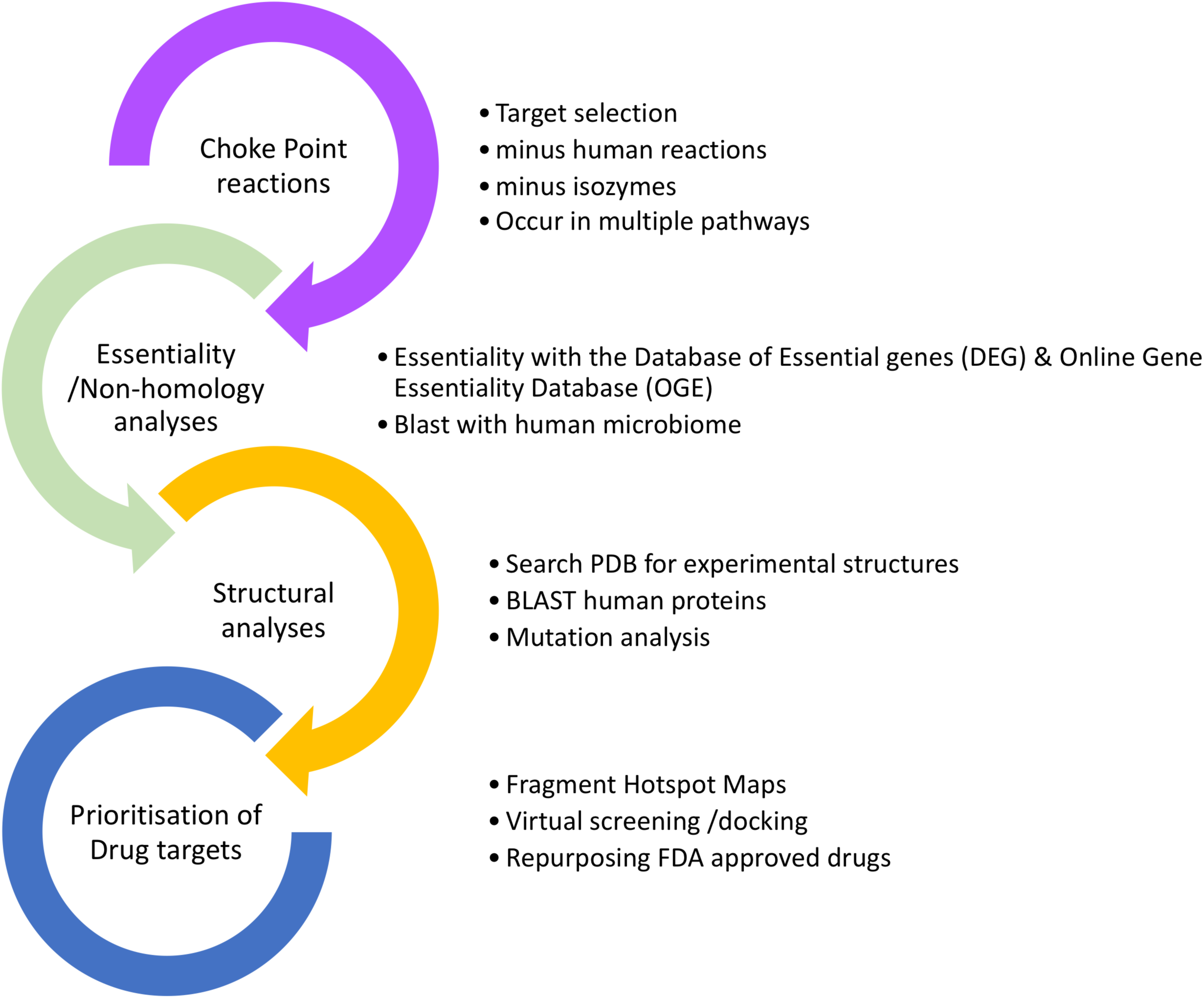
Pipeline for drug target identification in mycobacteria

### Chokepoint analysis for the identification of drug targets

The concept of ‘chokepoint reactions’ was introduced by Yeh and colleagues in 2004 (29), when they demonstrated how the metabolic network of an organism can be queried to predict potential drug targets. A chokepoint reaction of a metabolic network must not only be a reaction that is either the only consumer or the only producer of a given metabolite, but in addition, it is required that such a metabolite is not a dead-end metabolite, i.e. each chokepoint reaction should be balanced by at least one other reaction that produces or consumes that metabolite.

Chokepoint reactions are believed to be essential to the bacteria and are therefore potential drug targets. We initially analysed the three metabolic pathway databases of *M. tuberculosis, M. abscessus* and *M. leprae* for the unique reactions within the metabolic networks using the chokepoint algorithm. We excluded the chokepoint reactions found in humans and further restricted our selection to chokepoint enzymes that are present in more than one pathway in order to assess which enzymes will produce a greater impact on the metabolic network (Fig 2; S2 Table). Of the 333 potential drug targets initially identified in *M. tuberculosis*, 24 targets selected for further analyses were found to be involved in more than one pathway; these include lipid metabolism, coenzyme transport and metabolism, energy production and conversion, amino acid and nucleotide transport and metabolism (Table A in S2 Table). We postulate that potential inhibitors of targets identified in multiple pathways would have a more significant impact on the metabolic network of the bacterium than the inhibitors of a single pathway. In *M. abscessus*, 313 potential drug targets were initially identified and 15 were involved in more than one pathway, including cell wall and cell processes, coenzyme transport and metabolism, energy production and conversion, amino acid and nucleotide transport and metabolism; these targets were selected for further analyses (Table B in S2 Table). Similarly, 20 potential targets of the 183 identified in *M. leprae* were involved in more than one pathway including lipid transport and metabolism, nucleotide transport and metabolism, cell wall/membrane/envelope transport and metabolism/amino acid transport and metabolism / coenzyme transport and metabolism and these were selected for further analyses (Table C in S2 Table).

**Fig 2:**
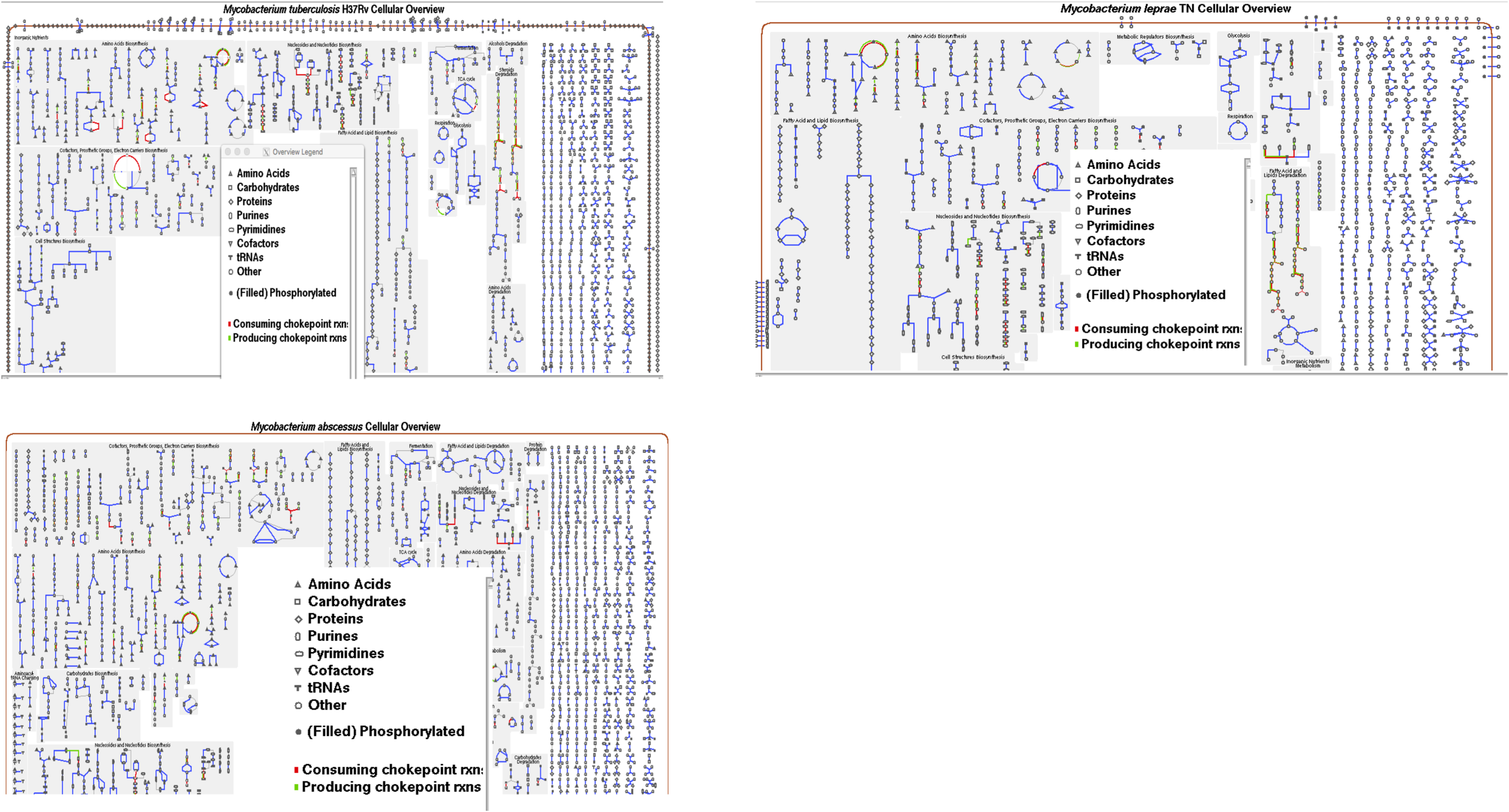
Choke point reactions in mycobacteria

Based on *in vivo* and *in vitro* essentiality patterns, we confirmed that 75% of the chokepoint enzymes are essential proteins to *Mycobacterium tuberculosis*. (33,34). Among the prioritized drug targets, 78% of the chokepoint enzymes are present in pathways shared between *M. tuberculosis, M. leprae* and *M. abscessus*; these include the biosynthesis of amino acids, such as arginine and ornithine, nucleotide transport and metabolism and pyrimidine biosynthesis (Tables A-C in S2 Tables). Here we focus on the arginine biosynthetic pathway in the *M. tuberculosis* network.

We performed Flux Balance Analysis (FBA) (35) on this pathway to assess the essentiality of ArgD, ArgH and other enzymes in the *M. tuberculosis* network. We implemented both FBA and Flux Variability Analysis (FVA) on the m7H10 medium (a growth medium of choice for cultivating M tb) using the ‘biomass’ reaction as the objective function and knocking out ArgD and ArgH to assess the impact on the metabolic network and to compute the minimum and maximum fluxes that the reactions can take. The maximum growth rates (h^-1^) predicted by FBA after knocking out ArgB, ArgC, ArgD and ArgH are reported in S3 Table.

A maximum growth rate of zero confirms the in-silico essentiality of the target enzymes ArgD and ArgH. Moreover, FVA (35) demonstrates a zero-flux distribution of the reactions when either of these essential enzymes is absent (Fig 3) in *M. tuberculosis*. Our FVA results also show that ArgF, ArgG and ArgJ exhibit zero flux distribution of reactions when they are absent from the pathway, although the growth rate of the bacterium when ArgA is knocked out is the same as the wild type. Therefore, we propose that ArgB, ArgC, ArgD, ArgF, ArgG, ArgH and ArgJ represent critical enzymes for the continued growth and viability of *M. tuberculosis* but ArgA is not essential to the bacterium.

**Fig 3:**
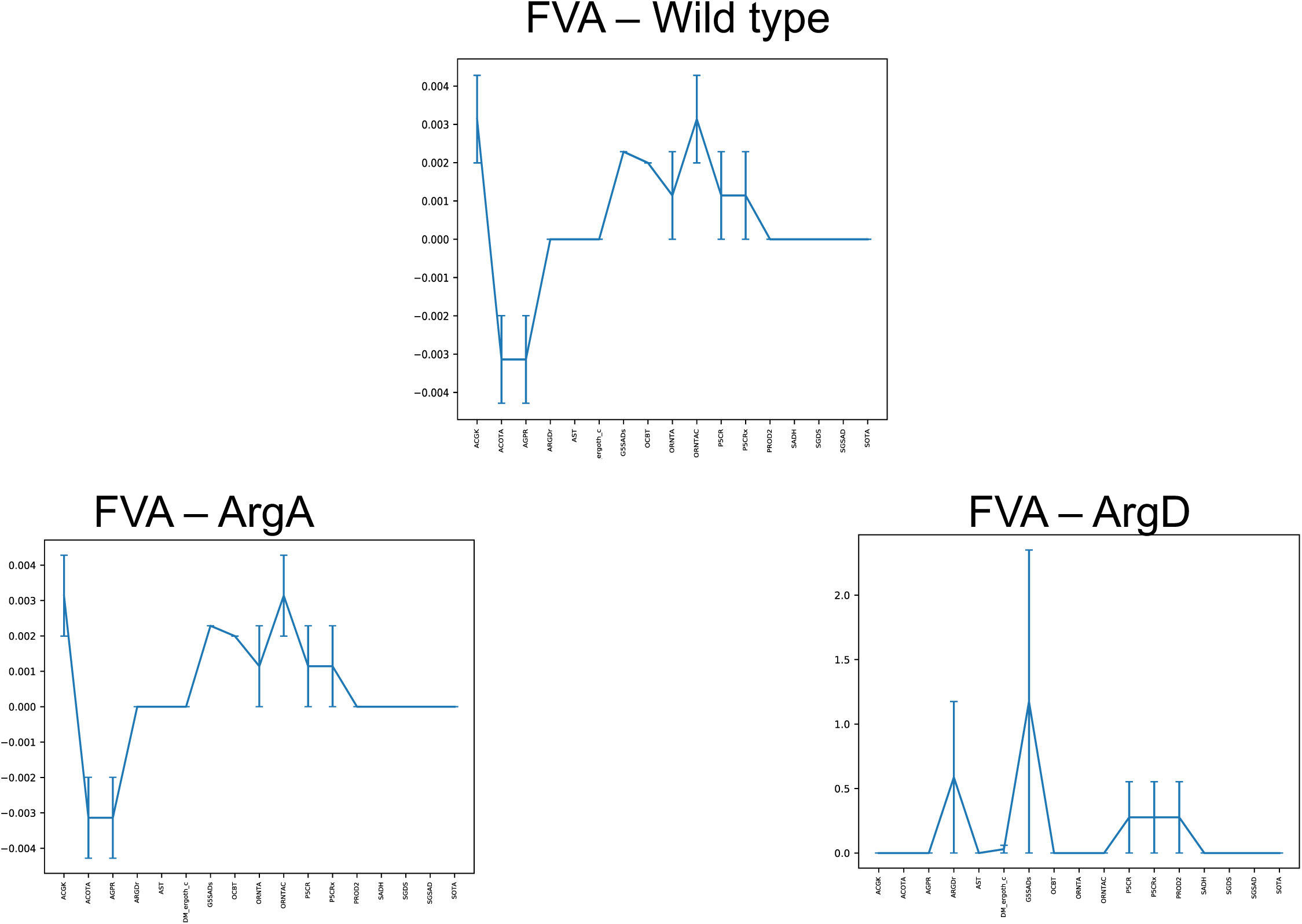
Flux Variability Analysis (FVA) demonstrates the difference in the flux of reactions in the arginine Biosynthetic pathway when enzymes, ArgA and ArgD are absent

### Comparative 3D modelling

We identified structures for ArgB, ArgC and ArgF, chokepoint enzymes present in *M. tuberculosis, M. abscessus*, and *M. leprae* that are present in the Protein Data Bank (PDB); these are listed on Tables A-C in S2 Tables. Comparative modelling techniques (36,37) were therefore employed to predict the structures for ArgD and ArgH that are not currently available in the PDB.

### ArgD

The ArgD (acetylornithine aminotransferase) protomer of *M. tuberculosis* was modelled using the templates 1vef (acetylornithine aminotransferase from *Thermus thermophiles*) with an identity of 45% and a coverage of 92% and 4adb (succinyl-ornithine transaminase from *E. coli* with an identity of 42% and a coverage of 96%). The overall Molprobity score of the model is 1.36 at the 98th percentile (38,39). Structural superimposition of the model with each of the templates 4adb and 1vef resulted in a root mean square deviation (RMSD) of 0.203Å and 0.908Å, respectively. A homodimer was then built by superimposing the protomers with each chain of the template 1vef; we have run Steepest Descent, Conjugate gradient and Modrefiner (40–42) to reduce steric clashes and to improve the quality of the models (Fig 4A in Fig 4).

**Fig 4.**
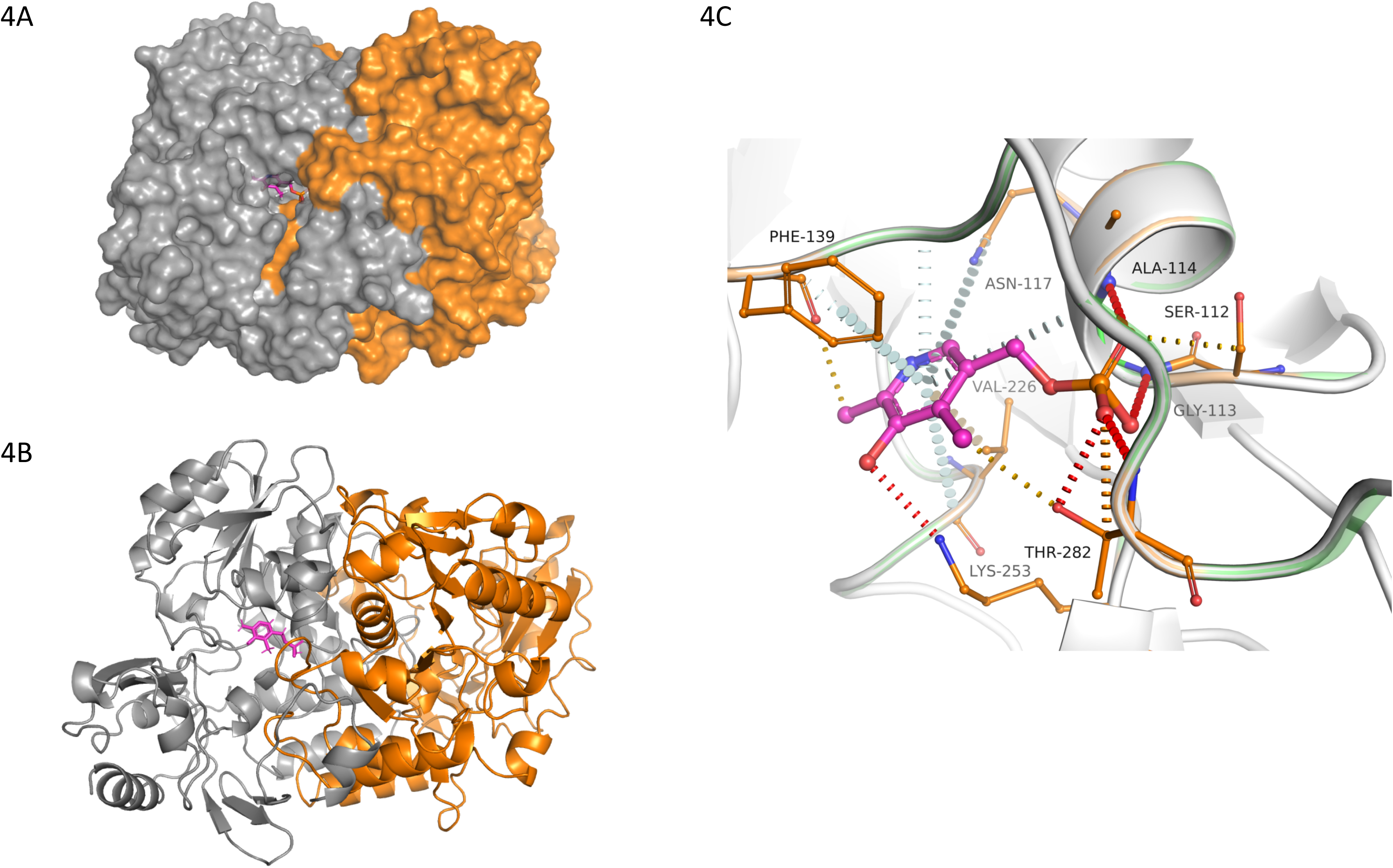
4A,B & C: Dimeric model of *Mtb* ArgD with PLP Binding site and interatomic interactions of PLP with the surrounding residue environment.

### ArgH

The ArgH (argininosuccinate lyase) protomer was modelled using the templates 2e9f (argininosuccinate lyase from *Thermus thermophilus* (Identity=45% and coverage=95%)) and 1k62 (argininosuccinate lyase from human (Identity=43% and coverage=97%)). The resulting model has an overall Molprobity score of 1.44 at 96th percentile (38,39). Structural superimposition with the templates 1k62 gave RMSD of 0.577Å and 1.069Å with 2e9f. A homotetramer was built by superimposing the protomer with each chain of the template 2e9f and we improved the quality of the models by running Steepest Descent, Conjugate gradient and Modrefiner (40–40) (Fig 5A in Fig 5).

**Fig 5:**
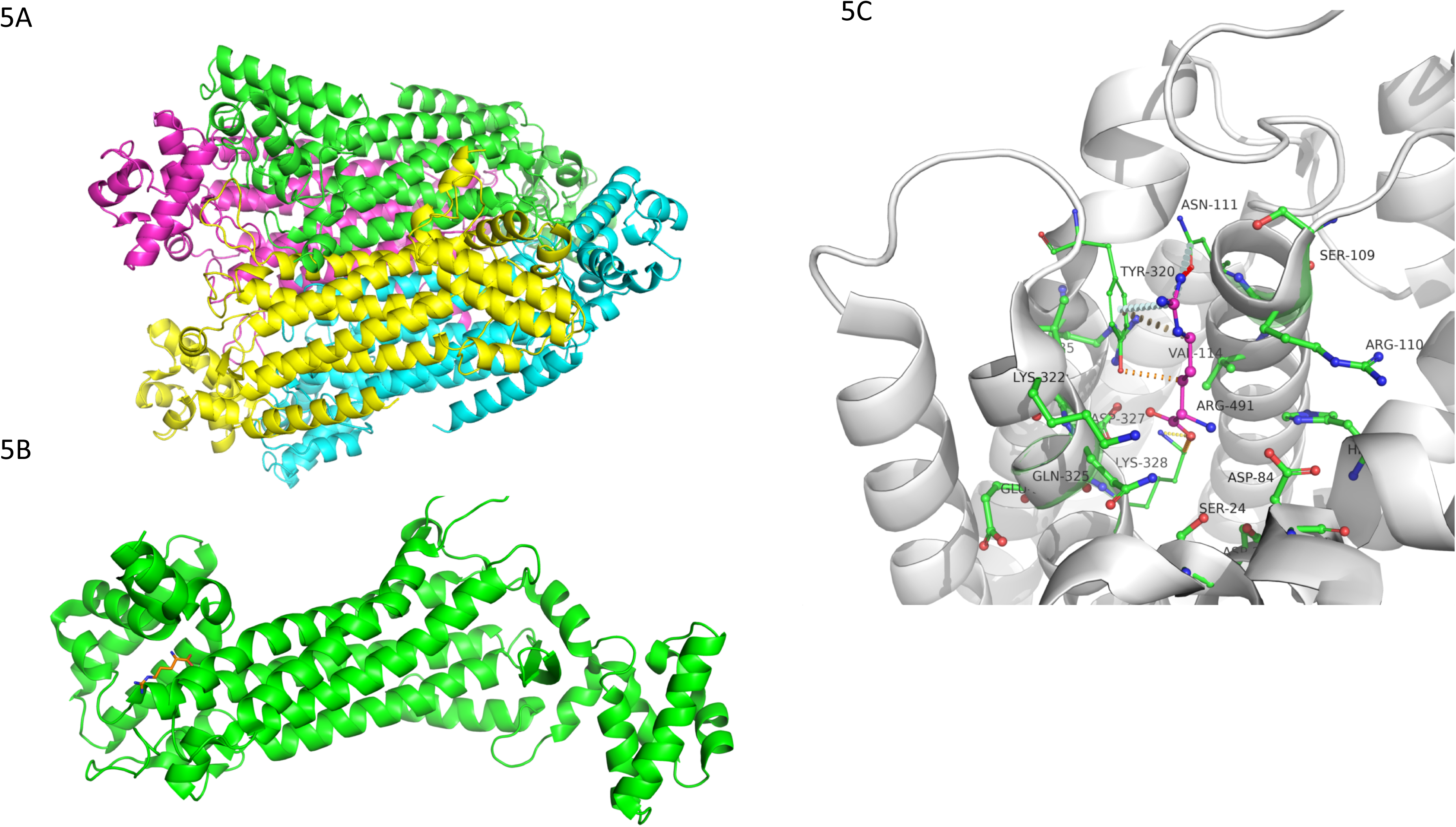
5A, B & C: Tetrameric model of *Mtb* ArgH with L-Arginine Binding site and interatomic interactions of L-Arginine with the surrounding residue environment.

In ArgD, the pyridoxal 5′-phosphate (PLP), the co-factor in the transamination reaction binding site, is located close to the dimeric interface (Fig 4B in Fig 4). It is located in the active centre cleft and forms multiple interatomic interactions with the surrounding residues. The oxygen atom of the phosphate moiety of PLP forms hydrogen-bond interactions with the main chain nitrogen atom of Thr282 (Fig 4C in Fig 4). It also forms weak polar interactions with side chain atoms of Thr282 and Val226. The pyridine ring forms multiple proximal π interactions with other the aromatic residues in the binding site. It forms a face to edge π interaction with Phe139 and carbon-π interaction with Val226. The Fragment Hotspots Maps which specifically highlight fragment-binding sites and their corresponding pharmacophores (43), overlapped the binding site corresponding to hydrophobic, donor and acceptor regions with a score of 17. ArgH, the L-arginine binding site is located at the interface between domains one and two (Fig 5B in Fig 5). Arginine interacts with the surrounding residues by the formation of a hydrogen bond with Asn111 and π interactions with Tyr320 (Fig 5C in Fig 5).

### Sequence-structure comparisons between human and Mtb ArgD and ArgH

A position-specific iterated BLAST (PSI-BLAST) search against the PDB of Mtb ArgD on the human proteome revealed six structural hits with sequence identities ranging from 30-36% (S4 Table). Alignment of *Mtb* ArgD sequence with the structure of 2BYJ (having the highest identity with the sequence) using JOY (44), revealed conserved residues across the PLP binding site between both the proteins. The alignments are depicted in Supp Fig 1A. Structural superimposition of chain A of ArgD model with chain A of 2BYJ resulted in an RMSD of 1.806Å (Supp Fig 1B).

PSI-BLAST of *Mtb* ArgH with the human proteome resulted in five hits on the PDB database with sequence identity ranging from 24-43% (S5 Table). L-Arginine in *Mtb* ArgH forms two hydrogen bonds, one with the sidechain oxygen atom of Asn111 and the other with sidechain nitrogen of the Lys328. It also forms hydrophobic contacts with Val114, Arg110 and Tyr320. JOY alignment (44) was performed for *Mtb* ArgH and 1AOS proteins and conserved regions were highlighted (Supp Fig 2A). Structural superimposition of chain A of human ArgD (1AOS) with chain A of Mtb ArgH model resulted in an RMSD of 0.751Å (Supp Fig 2B). In Mtb ArgD, three Fragment Hotspots were mapped in the binding sites for PLP, N-Acetyl-glutamate semialdehyde and glutamate (Supp Fig 3A) while in 2BYJ (Human ArgD), only one hotspot was mapped (Supp Fig 3B) at the PLP binding site that partially overlaps with that of the Mtb ArgD. Furthermore, in *Mtb* ArgH, three hotspots were mapped at the subunit interfaces (Supp Fig 4A). The maps were contoured at the score of 17 indicating higher likelihood for fragment binding. Conversely, no hotspots were mapped on the human ArgH (1AOS) at the score of 17 and few apolar regions were noted at a score of 14 (Supp Fig 4B). All the maps were at the subunit interfaces. Mishra and Surolia (45) have recently shown that ArgH undergoes conformational changes during catalysis that can influence the orientation of the substrate binding sites and subsequently the fragment binding.

The hotspots (43) and cavities (46) for ArgD were used to explore potential binding of two drugs, Amprenavir and Cochicine, to the ArgD protein and the putative interactions were investigated using a molecular docking approach. The structures of both ligands and of the pyridoxal 5’ phosphate (PLP) cofactor were retrieved from Drugbank in the simplified molecular-input line-entry system (SMILES) format (47). The SMILES strings were converted to the PDB format and their 3D structures were optimized after adding hydrogens using the *UFFOptimizeMolecule* function of RDKit. The resulting structures were used to generate pdbqt files, using the prepare_ligand.py script provided with MGLTools (48).

The modelled ArgD structure was converted to pdbqt format using MGLTools, and a cubic grid box with 24 Å-long edges was manually set to loosely accommodate the PLP binding site. Autodock VINA was used to generate up to 10 poses of each ligand (num_modes=10) within an energy range of 10 kcal/mol (energy_range=10) and the exhaustiveness was set to 40. The Open Drug Discovery Toolkit (ODDT) was used to re-score the docked poses, using the *RFScore_V3* function, trained on the *PdbBind2015* dataset. To increase the robustness of the results, the above-described procedure was repeated 200 times for each ligand and the results were clustered using the “gmx cluster” program, which integrates the GROMACS package (49). The clustering procedure was carried out with a RMSD cut-off of 0.2 nm and the docking poses were not fitted prior to the clustering, to capture translational and rotational differences. The resulting data were analysed using the DataWarrior software (50), which was also used to generate the scatter-plots and box-plots. The best-scoring pose of the cluster with the highest median score was selected and submitted to the PLIP web-server in order to analyse the formed interactions. The points of interaction of Amprenavir on the hotspot maps of the ArgD proteins on both human and M. tb are shown on Fig 6 and Supp Fig 3c. The PLP binding site identified on the ArgD sequence of M. tb is conserved in *M. leprae* and *M. abscessus*; the M. tb ArgD sequence is 83% identical to that of *M. leprae* and 74% identical to the *M. abscessus* ArgD sequence (Supp Fig 4). Similarly, the M. tb ArgH sequence is 88% identical to *M. leprae* and 79% identical to *M. abscessus*. The conserved domains of ArgD and ArgH indicate similar active and Fragment Hotspots in *M. leprae* and *M. abscessus* as seen in *M. tuberculosis* demonstrating the significance of metabolic pathway analysis in drug target identification of mycobacteria.

**Fig 6:**
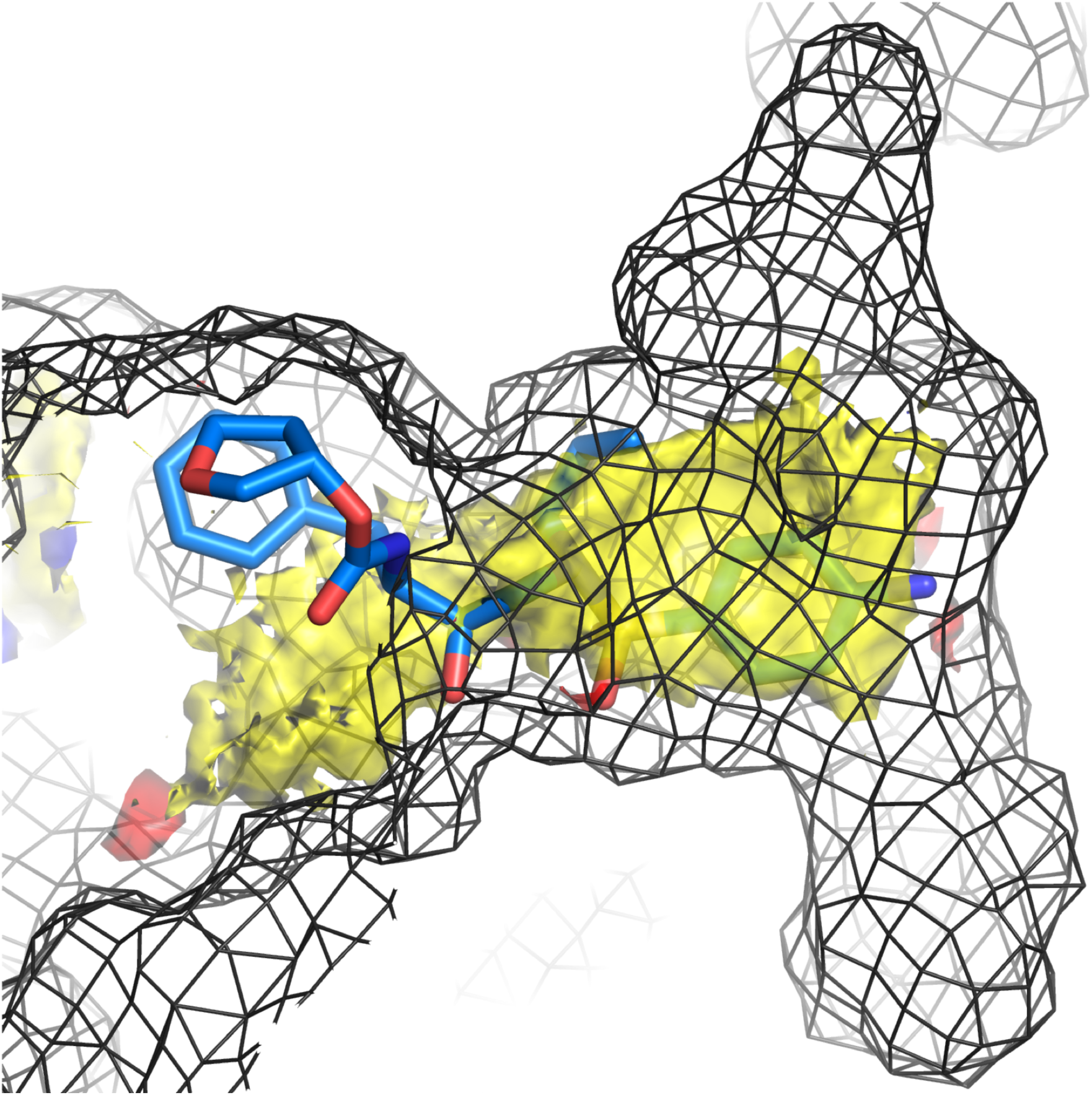
Superimposition of chosen docked pose of Amprenavir onto the predicted hot-spot map. The protein binding site volume is represented as dark-grey mesh.

## Discussion

The biosynthetic pathways identified as common between *M. tuberculosis, M. leprae* and *M. abscessus* include those for amino acids, carbohydrates, cell structures, aromatic compounds, cofactors, vitamin acids, fatty acids, metabolic regulators, secondary metabolites, as well as nucleoside and nucleotide biosynthetic pathways. These are all necessary for bacterial survival, growth, and division. We did not find the amine and polyamine biosynthetic pathways, which include the ectoine and glutathionylspermidine pathways in *M. leprae,* which were identified in *M. abscessus*; neither did we find the spermidine pathway in *M. leprae,* which is present in *M. tuberculosis*. The biosynthesis of ectoine enables *M abscessus*, an opportunistic bacterium, to survive in extreme osmotic conditions such as in soil and salt water environments. Also, the glutathionylspermidine biosynthetic pathway in *M. abscessus* and the spermidine biosynthetic pathway present in *M. tuberculosis* might be contributing to the maintenance of redox balance in *M. abscessus* and *M. tuberculosis*. As expected of an opportunistic organism such as *M. abscessus*, we identified the plant hormone biosynthetic pathway, indole-3-acetate biosynthesis, which is absent in the other two species, *M. tuberculosis* and *M. leprae,* which are obligate human pathogens.

Our studies on pathway analysis further demonstrate that there are more polysaccharide biosynthetic pathways in *M. tuberculosis* and *M. abscessus* than in *M. leprae*. This complements previous studies by (51), where they showed that other polysaccharides, in addition to cellulose, are key components in the biofilms of *M. tuberculosis*. Biofilm production, predominant in *M. tuberculosis, M. abscessus* and *Pseudomonas aeruginosa,* enables the bacteria to be more tolerant of antibiotics and persist in chronic disease. Species-specific reactions for *M. tuberculosis* that we identified include reactions in the cellulose biosynthetic pathway, although these reactions were missing in the most recent metabolic reconstruction of *M. tuberculosis* (30).

Common degradation pathways identified in the three species include include those for the turnover of alchohols, amino acids, carbohydrates, fatty acids, as well as those for nucleosides and nucleotides. Of particular note is the degradation of chlorinated compounds, such as dichlorobenzene, which have been observed in *M. abscessus* and *M. leprae* but not in *M. tuberculosis*. Previous studies (46, 47) have shown that this pathway is particularly exploited by *Pseudomonas spp*., which is able to use chlorinated compounds as sole carbon sources for nutrition, and this accounts for the prevalence of *Pseudomonas* in recreational waters. Therefore, we postulate that an opportunistic bacterium such as *M. abscessus* might be degrading dichlorobenzene as a carbon source whilst it is living in extreme conditions in the soil. We also hypothesize that the presence of a degradation pathway of dichlorobenzene in *M. leprae* suggests that the bacterium is able to survive in non-host environments by exploiting other carbon sources. This is consistent with previous studies (54,55), which showed that the presence of viable *M. leprae* in soil samples is suggestive of environmental reservoirs of the bacteria. The authors, therefore, provided an explanation for the occurrence of leprosy from individuals living in areas where human leprosy cases were not previously reported; this suggests that *M. leprae* might not really be an obligate parasite.

One of the characteristic features of *M. abscessus* is the large number of ABC and MmpL transporters proteins, ∼200, (56) that are dispersed in the genome (57) compared to that of 31 drug transporters in *M. tuberculosis* and 12 in *M. leprae* (58). The transporters are a very important set of proteins, involved in lipid, carbohydrate, nucleotide and amino-acid metabolism and enable *M. abscessus* to survive, replicate and maintain immunity within its host. In *M. leprae*, characterisation of the metabolic pathways is essential to provide insight into the vast array of unknown functions in the reduced genome. Including a comprehensive assembly of the transporter reactions in reconstructing the metabolic network of *M. abscessus* and updating the networks of *M. tuberculosis* and *M. leprae* will improve the *in-silico* predictions of essentiality and provide insight to the effect of drug uptake and resistance observed in *M. tuberculosis, M. abscessus* and *M. leprae*.

The above comparative studies using the metabolic pathway databases of *M. tuberculosis, M. abscessus* and *M. leprae* have allowed us to identify potential drug targets common to the three species using the chokepoint analysis and to demonstrate the effects of perturbations on the metabolic pathways. The identification of chokepoint reactions within the metabolic network can indicate the essentiality of genes required for the survival of the pathogen. Bringing chokepoint analysis of the reactions together with ligand-binding ability of the enzymes catalysing them is central to the selection of targets for drug discovery. We demonstrate the complementarity of these approaches using the arginine biosynthetic pathway as an example.

The following targets: ArgB, ArgC, ArgD and ArgH in the arginine biosynthetic pathway were among the prioritised drug targets identified through the chokepoint analysis in both *M. leprae* and *M. tuberculosis*. The four proteins occur in the full list of drug targets identified from the chokepoint analysis of *M. abscessus*. Furthermore, our FBA analysis shows that ArgB, ArgC, ArgD, ArgG, ArgH and ArgJ are all essential to the bacterium, ArgA amino-acid acetyltransferase, which catalyzes the production of N-acetyl-L-glutamate from L-glutamate is not essential to *M. tuberculosis* and absent in both *M. leprae* and *M. abscessus*. In *M*. *tuberculosis*, ArgJ can also play a dual function as it possesses the same acetyltransferase as ArgA and can also catalyse the production of L-ornithine from N-acetyl L-glutamate. Having considered all the proteins in the arginine biosynthetic pathway, we selected acetylornithine aminotransferase (ArgD) and argininosuccinate lyase (ArgH), currently with no experimental structures in the Protein Data Bank for further druggability and ligand-binding ability analysis including (i) sub-cellular localization (ii) check using similarity search in DrugBank (iii) comparative 3D-structure analysis of argD in mycobacterial species and humans and (v) pocket analysis (using Fragment Hotspot Maps).

Our analyses indicate that *M. tuberculosis* ArgD is cytoplasmic and also has significant homologues in DrugBank (with E-value < 0.00001). The druggable potential of ArgD and its applicability in re-purposing drugs was recently demonstrated on a previous study (46), for the identification of polypharmacological targets in *M*. *tuberculosis*, using a structural proteomic analysis of binding sites. The authors have comprehensively characterized the pocketome of *M. tuberculosis*, based on a structural proteomics strategy at genome-scale (46). This study indicates that ArgD in *M. tuberculosis* (Rv_1655) might be a target for drug-repurposing of compounds such as Amprenavir and Colchicine. Our analyses of pockets and hotspots show the Fragment Hotspot mapped around the phosphate moiety of the pyridoxal 5′-phosphate of the ArgD protein on *M. tuberculosis* and at the sub-unit interfaces on domain 2 of the *M. tuberculosis* ArgH protein.

A summary of interaction analysis carried out by PLP is shown in S6 Table and Supp Figure 7a depicts these interactions. Our results suggest that Amprenavir likely interacts with ArgD, whilst a strong interaction with Colchicine is less likely to take place (S6 Table). On the other hand, since Colchicine is a larger and less flexible molecule, docking assays that take receptor flexibility into account could prove to be more reliable. ArgH, on the other hand, demonstrates a promiscuity of interactions and fewer binding sites.

In a previous study, Tiwari and colleagues (59), have described ArgB and ArgJ as potential drug targets. On this study, we have evaluated all the enzymes on the arginine biosynthetic pathway, and provided:

a. The results of the chokepoint analyses (S2 Table) demonstrating that ArgB, ArgC, ArgD, ArgF, ArgH and ArgJ are potential drug targets in *M. tuberculosis, M. leprae* and *M. abscessus*.
b. We provide a list of pathways common to the three species, *M. tuberculosis, leprae and abscessus*, and specie-specific pathways for each mycobacterium (S1 Table).
c. The ligand-binding ability assessment of ArgD and ArgH (Fig 4 and 5) and comparative results with the arginine biosynthetic pathway enzymes in *M. leprae* and *M. abscessus*.
d. Comparative 3D-models of ArgD and ArgH and descriptions of their fragment hotspot maps.
e. Superimposition of chosen docked pose of Amprenavir onto the predicted hot-spot map of ArgD.
f. Python-based chokepoint algorithm for the drug target identification of a metabolic network file in SBML format and available for download (S1 Text)

Updated metabolic reconstructions of *M. leprae* and *M. abscessus* (Bannerman *et al*. manuscripts in preparation) will support future analysis for drug repurposing for the benefit of patients affected by infections of *M. tuberculosis, M. leprae* and *M. abscessus*.

## Methodology

### Pathway / Chokepoint analysis / Gene Essentiality

We performed a comparative analysis of pathways in the networks of *M. tuberculosis, M. abscessus* and *M. leprae* by using the most recent reconstruction of *Mycobacterium tuberculosis* metabolic network (30) and the automated build of the pathway genome database for *M. abscessus* and *M. leprae* using the Pathway Tools software (60). To determine potential drug targets, we implemented a pipeline (Fig 1), which includes an in-house chokepoint algorithm in Python, the input of which is a metabolic network expressed in the Systems Biology Markup Language (SBML) format, and the output is a spreadsheet with the reactions that are either the only producers or the only consumers of the metabolites. Metabolites are stored in different sheets according to the number of reactions that produce and consume them, e.g. all the metabolites that are produced by 2 reactions are contained in the same sheet. The sets of reactions that are the only producers, or the only consumers, of a given metabolite are also reported by the algorithm in the same spreadsheet. It should be noted that these sets of reactions can be seen as a generalization of the classical chokepoint algorithm, i.e. if the size of the set of reactions is one, then it corresponds to a chokepoint reaction, if the size is two, then the metabolite is either produced or consumed by at most two reactions, etc. The algorithm also offers the possibility of identifying the ‘dead-end’ metabolites, i.e. those for which there are no metabolic or transport reactions within the metabolic network, and which are not contained within the final biomass composition equations. Such metabolites indicate that the metabolic model is either incomplete or contains spurious pathways.

This algorithm was run on the *M. tuberculosis* iEKV1011 metabolic network model and compared with the BioCyc chokepoint implementation (29) on the models of *M*. *tuberculosis, M. abscessus* and *M. leprae*. We performed searches from literature, the database of essential genes (DEG) (33) and the Online Gene Essentiality (OGEE) (34) database to determine the *in vivo* and *in vitro* essentiality of the potential targets. We also performed Flux Balance Analysis (FBA) and Flux Variability Analysis (FVA) on the *M. tuberculosis* iEK1011 model (30) using the Python toolbox COBRApy (61).

### Structure modelling of selected drug targets

Comparative modelling of selected drug targets, ArgD and ArgH, chokepoint enzymes of the arginine biosynthetic pathway, which currently lack experimental structures in the PDB, was performed using an *in-house* modelling pipeline, *Vivace* (Ochoa M. *et al.*, Skwark M. and Blundell TL. manuscript in preparation). The pipeline was built using the Ruffus (Python) (Goodstadt, 2010) architecture that performs multiple tasks to generate models with substantial quality and reliability. In the modelling process, templates were selected from TOCCATA (Ochoa Montaño B, Blundell TL, unpublished) a database of consensus profiles built from CATH v4.2.0 and SCOPe v2.07 (62,63) based classification of protein structures (PDB files). PDBs within each profile are clustered based on sequence similarity using CD-HIT (64,65)) and structures are aligned using BATON, which was developed as a streamlined version of the in-house program COMPARER (66). Later FUGUE (63) is employed to generate target-template alignments from different profiles and profiles with the highest Z-scores are used in the downstream process. After further optimization of the clusters by discarding templates with more than 20% difference in sequence identity to the maximum hit, the remaining templates are classified into states based on ligand binding. Different states were generated for ligand-binding and one state for apomeric structure. Models are built for each of these states using MODELLER (36) version 9.19 (37).

Druggability checks of the selected targets, such as subcellular localization, were predicted using PSORTb a database of protein subcellular localization for bacteria and archaea (68), subCELlular LOcalization predictor, CELLO (69), TB-Pred (70). The identified target sequences were also searched against databases such as Drug Bank (target sequence collection for approved drugs) (71) and Therapeutic Target Database (TTB) (72), using default parameter settings of BLAST. The identification of significant hits in such databases is an indicator of the potential evidence of druggability of the target. AutoDock Vina was then used to investigate the potential interactions of suggested drug candidates with the proteins using a molecular docking approach (73).

## Acknowledgements

We thank the following funding agencies for providing support to the Blundell and Oliver groups. JJ: the Industrial Biotechnology Catalyst (Innovate UK, BBSRC, EPSRC) to support the translation, development and commercialisation of innovative Industrial Biotechnology processes; SCV: American Leprosy Mission; MJS: The Botnar Foundation (Project 6063); PT: The Cystic Fibrosis Trust (Strategic Centre Award-201); SM: the UK Medical Research Council (MRC-DBT Centre Partnership); AM: The Cambridge Commonwealth Trust (10380426) and the Pakistan HEC Cambridge Scholarship. The authors also thank Dr Bill Jacobs for useful discussions.

## Supporting Information

S1 Fig 1A. JOY Alignment of Human ArgD (2BYJ) with *Mtb* ArgD Model

S1 Fig 1B Superimposition of chain A of *Mtb* ArgD (Orange) with chain A of Human ArgD (Blue)

S2 Fig 2A. JOY Alignment of Human ArgH (1AOS) with *Mtb* ArgH Model

S2. Fig 2B. Superimposition of chain A of *Mtb* ArgH with chain A of Human ArgH (1AOS)

S3. Fig 3A. Three different hotspot maps in *Mtb* ArgD Model

S3 Fig 3B. One Hotspot map adjacent to the PLP binding site in Human ArgD (2BYJ)

S3 Fig 3c: Hotspot Maps on human ArgD 2BYJ and M tb ArgD

S4. Fig 4A. Three hotspot maps at the subunit interfaces in Mtb ArgH

S4. Fig 4B. Hotspot maps at the subunit interfaces in Human ArgH (1AOS)

S5. Fig 5A. Clustering of docking poses for the docked molecules -PLP

S5. Fig 5B. Clustering of docking poses for the docked molecules – Amprenavir

S5. Fig 5C. Clustering of docking poses for the docked molecules -Colchicine.

S6. Fig 6. Superimposition of chosen docked pose of Piridoxal 5’ Phosphate (in green) and the orientation obtained by X-ray crystallography (in cyan) after structural alignment of the experimental and modelled protein structures.

S1 Table. Summary of pathways present in *M. tuberculosis, M. abscessus* and *M. leprae* S2 Table. Chokepoint reactions present in *M. tuberculosis* (A), *M. abscessus* (B) and *M. leprae* (C)

S3 Table 3. Maximum growth rates (h-1) enzymes of the arginine biosynthetic pathway predicted by FBA

S4 Table 4. Sequence Structure comparisons between human and Mtb ArgD and ArgH S5 Table 5. PSI-BLAST of Mtb ArgH with human proteome

S6 Table 6. Interaction analysis carried out by PLP

S1 File. This is the Chokepoint algorithm

~~~
#! /usr/bin/env python #
# Algorithm to compute the chokepoint reactions, dead-ends, consumer and
# and producer reactions of metabolites.[Jorge Julvez and Bridget Bannerman]
#
# Input: Metabolic network in SBML formar or json format
# Output: Spreadsheet modelfile_chokeps.xls where modelfile is # the name of the file containing the metabolic network #
# Example of use: python chokeps.py --model mtb_sbml.xml
# In this case, the results are written to mtb_sbml_chokeps.xls
from__future__import division, print_function
import argparse
import cobra
import xlwt
def chokeps(modelfile):
  print("Reading the model…")
  if modelfile[-3:] == "xml":
    model = cobra.io.read_sbml_model(modelfile)
    xlsfile = modelfile[:-4]+'_chokeps.xls'
  elif modelfile[-4:] == "json":
    model = cobra.io.load_json_model(modelfile)
    xlsfile = modelfile[:-5]+'_chokeps.xls'
  else:
   raise RuntimeError("Model file must be either .xml .json")
  print("Model:", modelfile)
  print('Reactions:', len(model.reactions))
  print('Metabolites:', len(model.metabolites))
  print('Genes:', len(model.genes))
  for reac in model.reactions:
    if reac.lower_bound >= reac.upper_bound:
      print("Reaction:", reac.name, "has identical lower and
upper bound =", reac.lower_bound)
    elif reac.upper_bound <= 0.0:
      print("Reaction:", reac.name, "is backwards only")
  ###############
  chokedict = {-1: []} # -1 is the key of the dictionary for chokepoints
  for meta in model.metabolites:
  influxes, outfluxes = [], []
  for reac in meta.reactions:
   if reac.lower_bound >= 0.0: # forward reaction only
    if reac.metabolites[meta] > 0: # influx
      influxes.append(reac.id)
    elif reac.metabolites[meta] < 0: # outflux
      outfluxes.append(reac.id)
    elif reac.upper_bound <= 0.0: # backward reaction only
     if reac.metabolites[meta] > 0: # outflux
       outfluxes.append(reac.id)
     elif reac.metabolites[meta] < 0: # influx
         influxes.append(reac.id)
     else: # reversible reaction
       influxes.append(reac.id)
       outfluxes.append(reac.id)
    # Check if the metabolite is a chokepoint, i.e.
    #  (only one influx or only one outflux) and (is not a dead-end)
    #  and (influxes != outfluxes)
    if len(influxes) == 1 and len(outfluxes) >= 1 and influxes !
= outfluxes:
     chokedict[-1].append((influxes, meta.id, 'influx'))
   if len(outfluxes) == 1 and len(influxes) >= 1 and influxes !
= outfluxes:
     chokedict[-1].append((outfluxes, meta.id, 'outflux'))
     # Get the number of influxes and outfluxes of the metabolite
     if len(influxes) not in chokedict:
       chokedict[len(influxes)] = [(sorted(influxes), meta.id,
'influx')]
     else:
       chokedict[len(influxes)].append((sorted(influxes),
meta.id, 'influx'))
     if len(outfluxes) not in chokedict:
        chokedict[len(outfluxes)] = [(sorted(outfluxes),
meta.id, 'outflux')]
    else:
      chokedict[len(outfluxes)].append((sorted(outfluxes),
meta.id, 'outflux'))
print('Number of chokepoints with no dead-end metabolites:',
len(set([tup[0][0] for tup in chokedict[-1]])))
  print('Number of dead-end metabolites:', len(set([tup[1] for tup
in chokedict[0]])))
  # Write results to spreadsheet
  print("Writing results to", xlsfile)
  wb = xlwt.Workbook()
  bstyle = xlwt.easyxf('font: bold on')
  for choken in sorted(chokedict):
      if choken == -1:
         sheetname = "Chokepoints(no dead-ends)"
      elif choken == 0:
        sheetname = "Dead-ends"
      elif choken == 1:
       sheetname = "Chokepoints(includes dead-ends)"
      else:
      sheetname = str(choken)+" producers or "+str(choken)+"
consumers"
    ws = wb.add_sheet(sheetname)
    ws.write(0, 0, 'Reaction Id', bstyle)
    ws.write(0, 1, 'Metabolite Name', bstyle)
    ws.write(0, 2, 'Metabolite Id', bstyle)
    ws.write(0, 3, 'Type', bstyle)
    for row, tup in enumerate(sorted(chokedict[choken])):
      ws.write(row+1, 0, str(tup[0]))
      ws.write(row+1, 1,
model.metabolites.get_by_id(tup[1]).name)
      ws.write(row+1, 2, tup[1]) if tup[2] == "outflux":
      if choken == 0:
        msg = "No reaction consumes this metabolite"
      elif choken == 1:
        msg = "Only this reaction can consume this
metabolite"
      else:
        msg = "Only these reactions can consume this
metabolite"
      ws.write(row+1, 3, msg)
      else:
        if choken == 0:
          msg = "No reaction produces this metabolite"
        elif choken == 1:
          msg = "Only this reaction can produce this
metabolite"
      else:
          msg = "Only these reactions can produce this
      ws.write(row+1, 3, msg)
   wb.save(xlsfile)
return chokedict
# Parse arguments and call procedure
parser = argparse.ArgumentParser()
parser.add_argument("--model", dest='modelfile',
default='mtb_sbml.xml', help="File containing the model")
args = parser.parse_args()
chokeps(args.modelfile)
~~~

